# Temperature and photoperiod differentially impact maternal phenotypes in diapause egg-laying *Aedes albopictus* mosquitoes

**DOI:** 10.1101/2024.04.19.590231

**Authors:** In Hae Lee, Anthony S. Lee, Helen V. Kogan, Linhan Dong, Laura B. Duvall

## Abstract

**Background:** *Aedes albopictus* (Skuse 1894) mosquitoes can transmit deadly arboviruses and are globally invasive due to their ability to survive in both tropical and temperate climates. Although adults cannot survive harsh winters, females are capable of anticipating seasonal change and producing overwintering diapause (DP) eggs that remain in a state of arrested development over the winter and hatch when favorable conditions return in the spring. Previous work has shown that shortening photoperiod (day length) alone is sufficient for DP induction. While decreasing temperatures can facilitate DP entry, temperature signals alone are not sufficient to induce DP.

**Methodology/Principal Findings:** To identify maternal phenotypes predictive of DP egg production, we characterized aspects of maternal physiology and behavior to identify those that correlate with DP egg production and changes in photoperiod, versus changes in temperature. Neither changes in temperature nor photoperiod impacted protein preference, blood meal consumption, or total number of eggs produced per female. Egg retention and oviposition timing were influenced by temperature, independent of DP egg production. However, females housed under short photoperiod conditions showed increased starvation resistance, despite showing similar levels of locomotor activity and internal stores of triacylglycerols, glucose, glycogen, and trehalose compared to females housed in long photoperiods.

**Conclusions/Significance:** These results suggest that temperature and photoperiod differentially affect maternal phenotypes and identify starvation resistance as a maternal phenotype that is influenced by photoperiod and can be used to predict DP egg status.

**AUTHOR SUMMARY:** *Aedes albopictus* mosquitoes can survive in temperate climates because females are able to detect changes in temperature and day length in the autumn to produce “diapause” eggs that are provisioned with extra nutrients to survive harsh winters by remaining in a state of arrest and hatching in the spring. Although temperature and day length normally change in concert with each other throughout the year, studies have shown that day length is the most important cue used by female mosquitoes to initiate diapause egg production. In this study, the authors examined aspects of feeding and reproduction to determine which of these correlated with day length. Many aspects of reproduction were influenced by temperature; however, housing female mosquitoes under autumn-like short day conditions led to increased starvation resistance. Surprisingly these females survive longer even though they do not appear to expend less energy compared to females housed in spring-like long day conditions at the same temperature. This suggests that the mosquito mothers undergo changes in their metabolism that can be used predict whether she will produce eggs that can survive the winter. This new knowledge may lead to new targets to disrupt seasonal reproduction in mosquitoes and limit the parts of the world where they can survive.

## INTRODUCTION

The Asian tiger mosquito, *Aedes albopictus*, is one of the world’s most dangerous invasive species (1). Female *Ae. albopictus* are obligate blood-feeders that require host blood protein to complete reproduction and can transmit pathogens to humans that cause diseases including yellow fever, chikungunya, dengue, and Zika (2–7). Its ancestral range likely spanned from India to Japan but this species has successfully invaded substantial portion of the globe, including North America, and climate change modeling predicts that these animals will continue to expand and redistribute their geographical range, increasing net and new exposure to *Aedes*-borne diseases (1,8,9). One key to their ability to invade diverse ecological spaces is the capacity of females to detect seasonal changes and produce developmentally arrested diapause (DP) eggs (10–12). In temperate climates, *Ae. albopictus* adult females sense and anticipate seasonal changes to produce DP eggs that survive winter and hatch the following spring (10,11,13). Maternally-instigated embryonic DP maximizes progeny survival, ensuring overwintering survival of the species (9). Although DP induction is found in many insect species, the stage of arrest and mechanisms of induction are varied: in the silkworm, *Bombyx mori,* the neuropeptide diapause hormone (DH) acts on the mother’s ovaries to influence carbohydrate metabolism, which in turn influences the DP fate of her eggs, but DH is not to be present in Diptera (14). Traditionally, DP egg status is scored by performing hatch assays over days or weeks (10,15,17). In *Ae. albopictus*, DP eggs contain higher total lipid levels compared to non-DP eggs and previous work has identified transcriptional changes that suggest that females alter lipid metabolism and energy expenditures in preparation for the production of DP eggs (15,16) suggesting that maternal metabolic phenotypes may be earlier predictors of future DP egg status.

Previous work has shown that shortening photoperiod alone is sufficient for DP induction (18,19). While decreasing temperatures can facilitate DP entry, temperature signals alone are not sufficient to induce DP (20). The aim of this study was to characterize maternal feeding, reproductive, and metabolic phenotypes to identify those that correlate with photoperiod versus those that correlate with temperature. While temperature influenced multiple features of reproduction, we found that photoperiod influenced female starvation resistance. Surprisingly, females housed in low temperature and short photoperiod showed increased starvation resistance, even though they maintained similar levels of locomotor activity and nutrient utilization compared to females reared under long photoperiod conditions at the same temperature. This indicates that females undergo energetic changes that precede DP egg production and can inform future experiments to identify the specific metabolic changes that underlie photoperiod-associated starvation resistance.

## RESULTS

### Diapause egg production is induced by short photoperiod

We used established environmental conditions to induce DP egg production (10,15,17,24–26). Eggs from the same clutch were common-garden reared under high temperature, long photoperiod conditions (26°C, 16L:8D) to ensure that larval nutrition and developmental timeline were identical for all groups. At the pupal stage, when feeding was complete, animals were moved into one of three environmental conditions: high temperature, long photoperiod (26°C, 16L:8D); low temperature, long photoperiod (21°C, 16L:8D); or low temperature, short photoperiod (21°C, 8L:16D) (Figure 1A). Adults in each condition were allowed to mate freely and maximal mating rates were achieved for all groups within 3 days (Supplemental Table 1).

**Figure 1.**
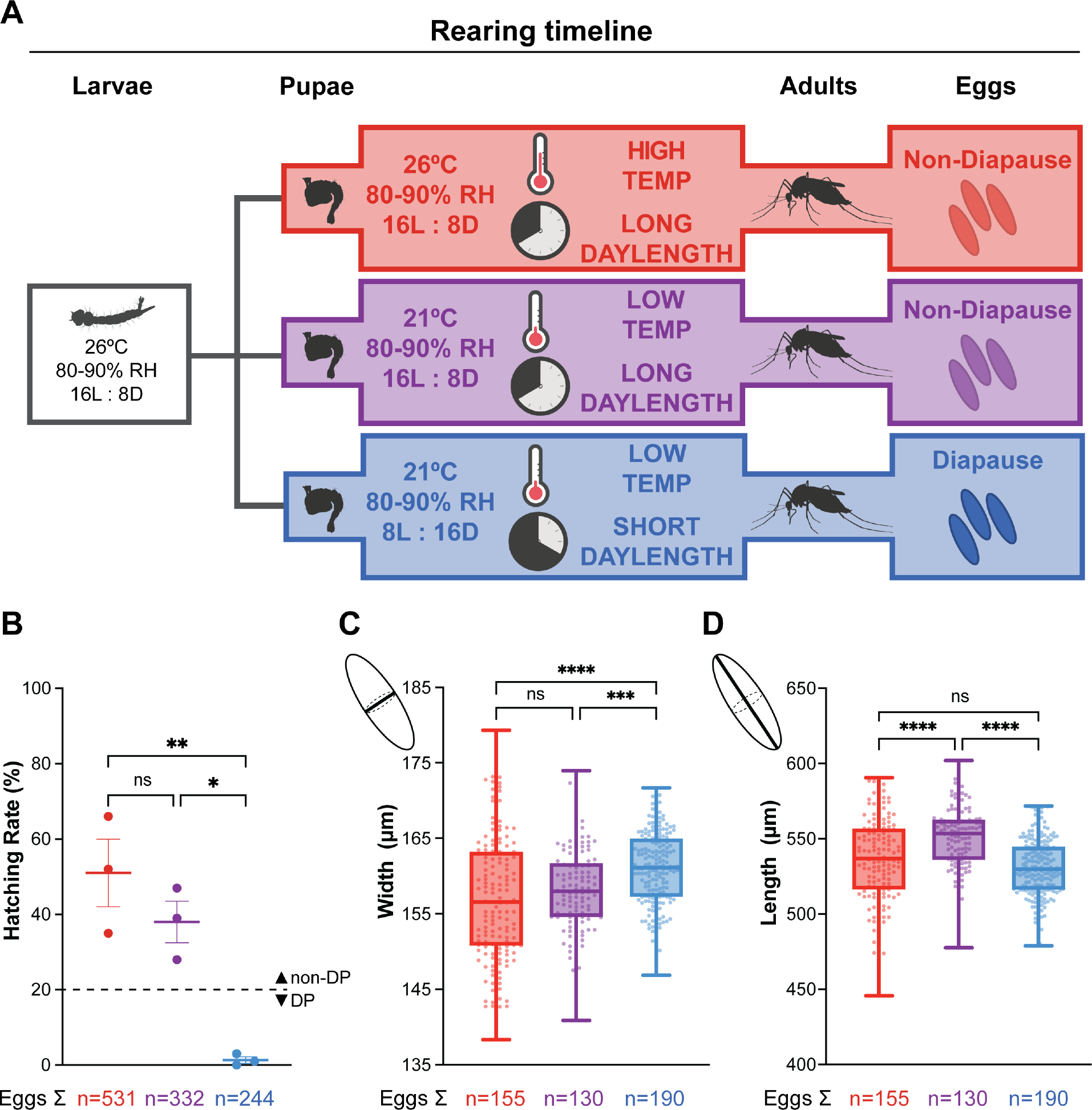
Short photoperiod induces diapause egg production (A) Schematic of rearing conditions, developmental timeline, and diapause (DP) egg status. Eggs from the same clutch were common-garden reared under 26°C, 16L:8D conditions. At the pupal stage, animals were separated into three conditions: high temperature, long photoperiod (red); low temperature, long photoperiod (purple); low temperature, short photoperiod (blue). B) Hatching assay to determine DP egg status. % of eggs hatched after exposure to hatching stimulus within 3 weeks of oviposition. Data is shown as mean with SEM (n = 3 replicates; >50 eggs per replicate; one-way ANOVA followed by Tukey’s multiple comparisons test; * = p < 0.05, ** = p < 0.01). C and D) Egg width (C) and length (D) measurement. Data is shown as median with range (n = 2 replicates; 130 - 190 eggs per replicate; one- way ANOVA followed by Tukey’s multiple comparisons test; *** = p < 0.0005; **** = p < 0.0001).

Instead of hatching immediately, DP eggs remain developmentally arrested for 2-3 months to survive winter. To confirm DP egg status, we used an established hatching assay and measured egg sizes (10,15,17,24–26). Consistent with previous reports, eggs laid by females housed under long photoperiod conditions at either 26°C and 21°C readily hatched, while eggs laid by females housed under short photoperiod conditions at 21°C were embryonated and viable but did not hatch in response to the same stimulus within 3 weeks of oviposition (Figure 1B). We measured the width (Figure 1C) and length (Figure 1D) of eggs laid by females housed in these 3 different conditions. Eggs laid by females housed in low temperature, short photoperiod conditions were significantly wider (160.9 ± 0.36 μm) than those laid by females housed under long photoperiod conditions at either 26°C (157.0 ± 0.67 μm) or 21°C, (158.1 ± 0.51 μm) consistent with previous studies (10,15,24–26). Altogether, females housed at 21°C laid DP eggs only under short photoperiod but not long photoperiod conditions. These findings recapitulate previous studies (10,15,24–26) and demonstrate that short photoperiod is the primary driver of *Ae. albopictus* DP egg production.

### Temperature impacts meal size, but not protein preference, in a two-choice assay

Temperature and photoperiod can influence nutrient preferences in diapausing insects; diapausing *Drosophila* change their preferred food from protein-rich yeast to plants (27). In *Culex pipiens*, a mosquito species in which DP occurs in adult females, the diapausing females suppress blood feeding and increase their sugar consumption (28,29). We therefore asked whether maternal meal consumption or preference change in *Ae. albopictus* females housed under the conditions depicted in Figure 1A. Although nutrients from sucrose are sufficient for survival, females require blood protein to develop their eggs. Previous work has shown that DP eggs are nutrient-rich with 30% higher lipid content compared to non-DP eggs (16,30,31). We reasoned that females producing DP eggs may show altered nutrient preferences to effectively provision these nutrient-dense eggs. Females employ two distinct feeding programs when ingesting blood versus sugar meals. Once meal types are distinguished by the feeding appendages used to evaluate tastants, females ingest starkly different sized meals when ingesting blood versus sugar. Females reliably double their body weight when consuming blood meals, whereas females take smaller sugar meals (32). The fate of these nutrients differ as well, with blood being allocated mostly for reproduction and sugar being allocated for metabolism (32–36). To ask if DP egg-producing females prefer a protein-rich meal, we compared female’s preference for sugar meals (10% sucrose) versus sugar meal with added protein (10% sucrose + 10% BSA; “protein rich meal”) in a two-choice assay. These two meals are consumed using the same sugar feeding program, allowing us to make direct comparisons between meal sizes. Females were allowed to feed *ad libitum* for 24 hours with access to both meals that can be distinguished by fluorescent labels. Females were collected at the end of the assay and scored individually for fluorescence intensity to calculate the volume of each meal consumed.

Females from all three groups showed a similar preference for the sucrose + BSA meal (high temperature, long photoperiod 17.91 ± 7.1%; low temperature, long photoperiod 13.93 ± 13.32%; low temperature, short photoperiod 26.27 ± 12.02%) (Figure 2A). Interestingly, females housed in high temperature, long photoperiod conditions consumed smaller meals overall (0.448 ± 0.060 μl) compared to females housed in low temperature conditions in both long (0.968 ± 0.107 μl) and short (1.03 ± 0.0802 μl) photoperiods (Figures 2B). Although DP egg-producing females housed in the low temperature, short photoperiod condition consumed larger volumes of both meals compared to females housed in high temperature, long photoperiod conditions, meal sizes were not significantly different from the meals consumed by non-DP producing females housed under low temperatures and long photoperiod conditions (Figures 2C and 2D). Meal preference was unaffected by environmental manipulations and differences in the combined meal size were driven by changes in temperature and not predictive of DP egg status.

**Figure 2.**
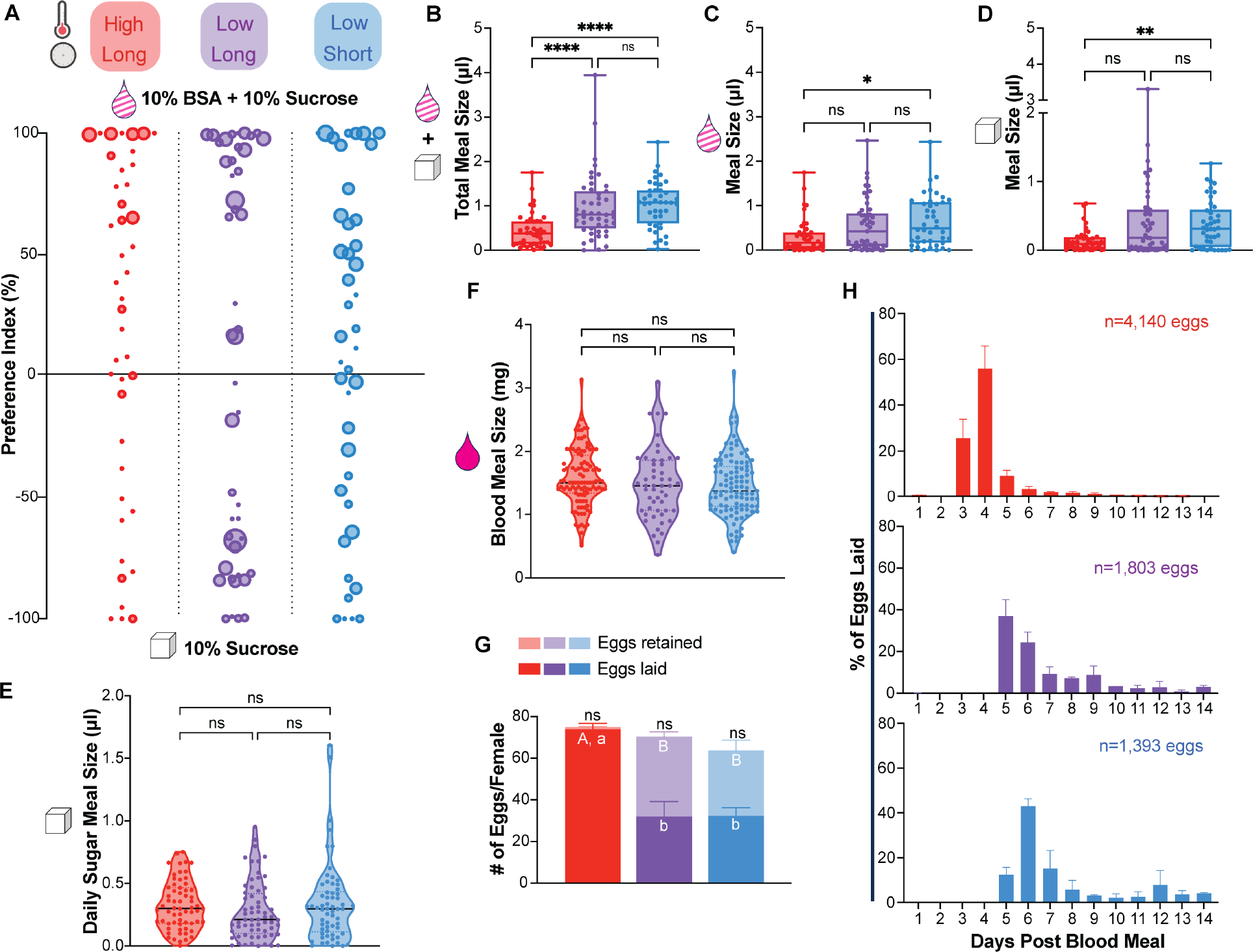
Temperature impacts meal size in a 2 choice assay and egg retention. A) Two-choice assay to measure preference for “no protein” sugar meal (10% sucrose; bottom) versus “protein rich” meal (10% sucrose + 10% BSA; top). Dot size represents total meal size. n = 6 replicates, 4 – 14 females per round. B-D) Meal size for B) combined “sucrose” and “BSA + sucrose” meals; C) “BSA + sucrose” meal only; D) “sucrose” meal only. Data is shown as median with range (n = 6 replicates, 4 – 14 females per replicate; Kruskal Wallace with Dunn’s multiple comparisons test; * = p < 0.05, ** = p < 0.001, *** = p < 0.001, **** = p < 0.00001). E) Sugar meal size in a “no choice” assay. Data is shown as median with range (n = 4 - 5 replicates, 7 - 16 females per replicate; Kruskal Wallace with Dunn’s multiple comparisons test, n.s. = p > 0.05. F) Quantification of blood meal size in “no choice” assay. Data is shown as median with range (n = 2 - 5 rounds, 13 – 28 females per round; Kruskal Wallace with Dunn’s multiple comparisons test; n.s. = p > 0.05). G) Number of eggs produced per female including eggs laid and retained in the ovary. Data is shown as mean with SEM (n= 2 - 5 replicates; n = 13-28 females per replicate; one-way ANOVA followed by Tukey’s multiple comparisons test, n.s. = p > 0.05; letters indicate statistically distinguishable groups (p < 0.001); capital letters indicate eggs retained and lower case letters indicate eggs laid). H) Daily oviposition measured for days 1-14 post blood meal (n = 2 replicates; 21 - 28 females per replicate).

### Neither temperature nor photoperiod impacts sugar or blood feeding alone

Nectar and blood meals are normally consumed via distinct mechanisms and blood meals are significantly larger than sucrose meals (32,37). Therefore, we next asked whether baseline sugar or blood feeding were impacted by temperature or photoperiod. In no-choice sugar feeding assays, females were offered a single fluorescently-labeled meal of 10% sucrose and consumption was quantified each day for 4 consecutive days after eclosion. Females from all three groups consumed similar amounts of sucrose (Figure 2E). To quantify blood feeding, females were offered sheep blood in an artificial membrane feeder and blood meal size was determined by weighing females before and after feeding. Females consumed similar sized blood meals in all three conditions.

Altogether, changes in temperature and photoperiod did not impact the amount of sugar or blood consumed alone (Figure 2E-F).

### Low temperature delays oviposition and induces egg retention

Protein from blood nutrients is required for egg production and development. To evaluate whether changes in temperature and photoperiod impact egg production, females were blood fed individually and allowed to lay eggs for two weeks, at which point we counted the total number of eggs produced by each female. In all conditions, females produced the same total number of eggs; however, there were differences in the number of eggs laid versus retained in the ovary after two weeks (Figure 2G). Females housed under high temperature, long photoperiod conditions laid the majority of their eggs (73.842 ± 2.915 eggs laid versus 1.074 ± 0.377 eggs retained); however, females housed in low temperature conditions retained roughly half of their eggs under both long (31.935 ± 7.769 eggs laid vs 38.417 ± 1.750 eggs retained) and short photoperiod conditions (32.318 ± 3.939 eggs laid vs 31.366 ± 4.936 eggs retained). To confirm that retained eggs were still viable, we allowed females to lay their remaining eggs until death and confirmed egg embryonation (17) (Supplemental Figure 1). We also analyzed temporal aspects of egg-laying and found that females housed in both low temperature conditions showed a 2-day delay in oviposition compared to females under high temperature, long photoperiod conditions (Figure 2H). These data show that females housed in low temperatures retained eggs and delayed oviposition, but these changes were not predictive of DP egg status.

### High temperature increases oocyte heterogeneity before blood consumption

Based on the delayed oviposition phenotype we observed (Figure 2H), we predicted that oocyte development would also be delayed in females housed under low temperature conditions. We dissected ovaries from females housed under all three conditions at timepoints before blood feeding and at 1, 3, 7, and 14 days after a blood meal and staged ovarian development by calculating the percent of the ovariole occupied by the nurse cells (Figure 3). In immature ovarioles, the nurse cells occupy ≥ 75% of the total ovariole area but occupancy decreases as eggs develop until the nurse cells are no longer present in mature eggs (38,39). *Ae. albopictus* females are obligate blood feeders and it is generally thought that ovarioles are maintained in a state of previtellogenic arrest, which is released after the female takes a blood meal (40). Consistent with this prediction, we found that the ovaries of females housed in low temperature conditions contained only immature and early stage ovarioles, regardless of photoperiod. Surprisingly, females housed in high temperature, long photoperiod conditions showed a heterogenous mix of oocyte developmental stages before blood feeding, including some mature eggs (Figures 3A-B). After blood feeding, ovary development became synchronous across all 3 conditions with mature eggs developing by day 3 in all groups (Figure 3A-B). Females housed in high temperature, long photoperiod conditions laid most of their eggs by day 7, consistent with Figure 2H, and had “reset” to produce new early stage ovarioles by day 14. However, females housed in low temperature under both long and short photoperiod conditions maintained a heterogenous mix of oocytes and mature eggs even at 14 days after blood feeding, consistent with their delayed oviposition timeline (Figure 2H, 3B). Overall, high temperature increased oocyte heterogeneity before blood feeding but did not impact egg development after blood feeding and changes in photoperiod did not impact oocyte development.

**Figure 3.**
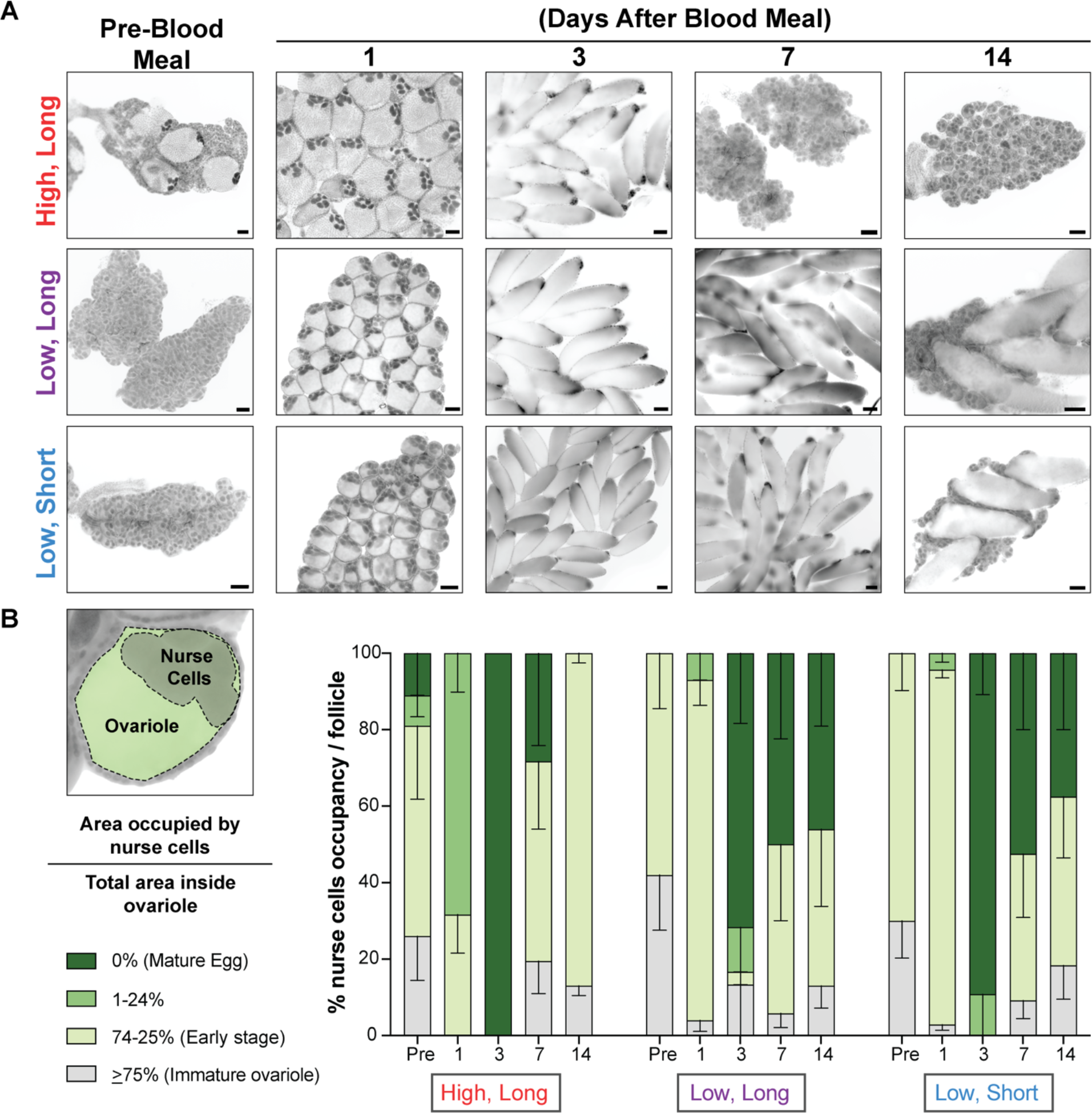
Temperature impacts oocyte heterogeneity before blood feeding. A) Representative images of ovarioles from females at pre-blood meal, and 1, 3, 7, and 14 days after a blood meal housed in the conditions indicated. B) Left diagram shows an example of a primary ovarian follicle. Ovarian development was measured by calculating the percentage of the follicle (green) occupied by the nurse cells (gray). As the follicle matures into an egg, the percentage occupied by the nurse cells decreases. Right graph shows all oocyte developmental stages at each timepoint and environmental conditions. Each timepoint includes measurements from 20-30 ovarioles from 3-7 pairs of ovaries. Data is shown as mean with SEM.

### Short photoperiod increases starvation resistance

So far, we examined the effect of nutrients obtained from external blood and sucrose meals. Previous work suggests that animals can dramatically alter their metabolic states to support DP (16,29,41) so to investigate whether environmental conditions impact the female’s ability to utilize internal nutrient stores, we tested maternal starvation resistance. Newly-eclosed females were given access to water only and checked daily for mortality. Although metabolic changes are often primarily associated with temperature (42), we unexpectedly found that photoperiod, but not temperature, impacted survival. Females housed in low temperature, short photoperiod conditions survived 2-3 days longer (median survival: 9 days) compared to females housed in long photoperiod conditions in both high (median survival: 6 days) and low temperatures (median survival: 6 days) (Figure 4A). Larval nutrition cannot account for these differences as all groups were common garden reared during development (Figure 1A). Starvation resistance could be accounted for by differences in daily energy expenditure or levels of internal nutrient stores in adult females. To ask whether environmental conditions altered locomotor activity during periods of starvation, we recorded daily locomotor activity of individual females using video tracking in a custom-built arena for 3 days following eclosion. Total locomotor activity was highest for animals housed in the high temperature, long photoperiod conditions (72.89 ± 17.64 cm) compared to animals housed in low temperature conditions in both photoperiods (long photoperiod: 47.55 ± 26.21 cm; short photoperiod: 34.72 ± 17.12 cm) (Figure 4B). Photoperiod change alone did not alter total locomotor activity and is not sufficient to account for differences in starvation resistance.

**Figure 4.**
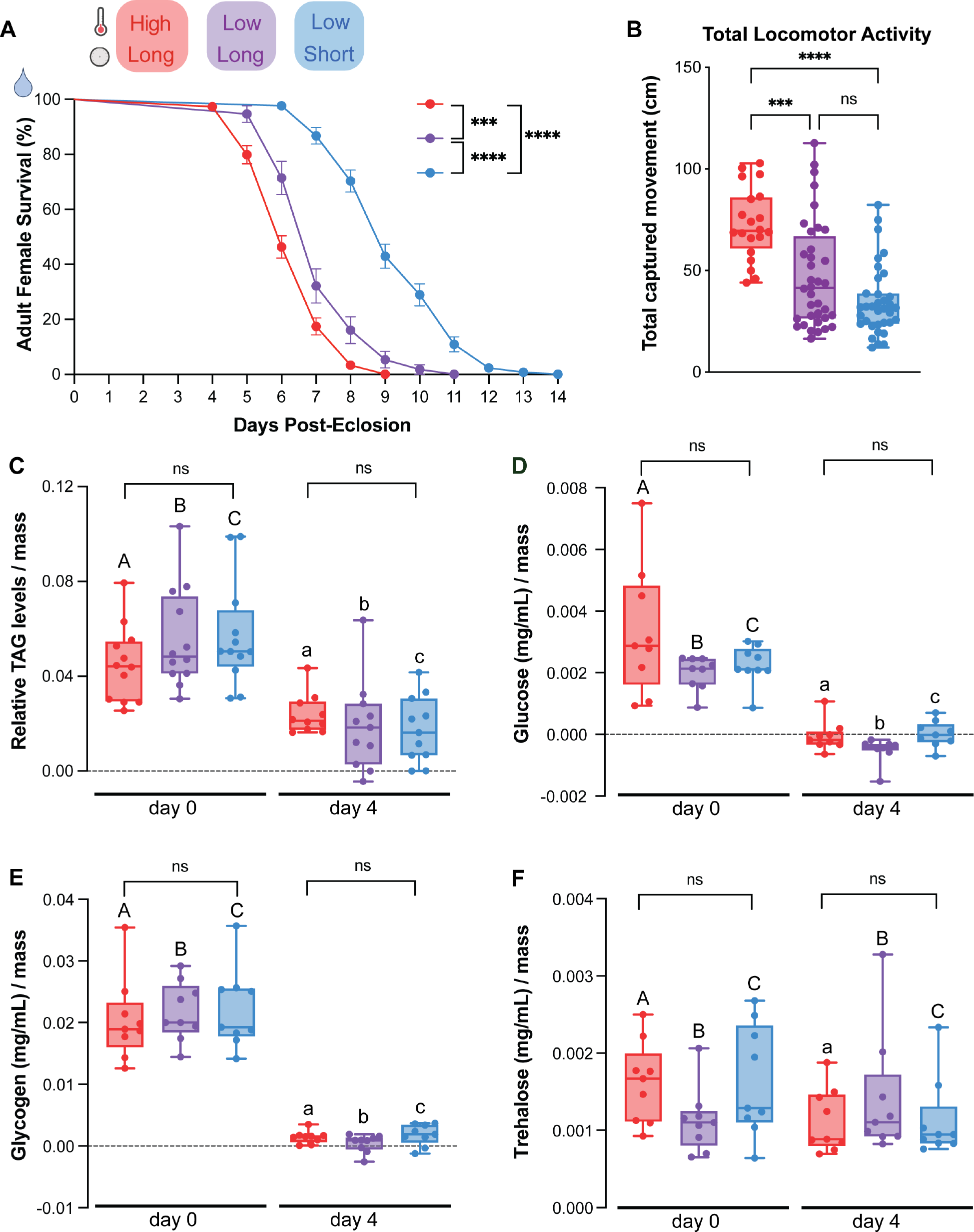
Short photoperiod increases starvation resistance. A) Daily survival for adult females provided with access to water only. Data is shown as mean with SEM (n = 2 - 4 replicates, 24 – 42 females per replicate; Log-rank (Mantel-Cox) test; *** = p < 0.0005, **** = p < 0.00001). B) Total daily locomotor activity over 3 days by females provided access to water only. Data is shown as median with range. (n = 3 replicates, n = 6 – 16 females/replicate; Kruskal Wallace with Dunn’s multiple comparisons test; *** = p < 0.0005, **** = p < 0.00001) C - F) Relative C) triacylglycerols (TAG), D) glucose, E) glycogen, and F) trehalose levels in whole females (groups of 5) collected after 0 and 4 days of starvation. Data is shown as median with range. n = 3 replicates, 4 groups of 5 females per replicate; Kruskal Wallace with Dunn’s multiple comparisons test was performed to compare all groups at day 0 and day 4; n.s. = p > 0.05; Mann-Whitney Test was used to compare matched pairs of conditions between day 0 and day 4; letters indicate matched pairs and upper vs lower letter case indicates statistically distinct groups p < 0.05).

### Starvation resistance is not accounted for by utilization of TAGs, glucose, glycogen, or **trehalose**

To ask whether utilization of internal nutrient stores can account for starvation resistance, we investigated triacyclglycerols (TAGs), a major component of lipid energy storage, as well as glucose, glycogen, and trehalose. Based on the survival curves in Figure 4A, we chose to quantify these nutrients on day 0 and after 4 days of starvation, a timepoint in which we saw low levels of lethality for all groups, shortly preceding the onset of lethality in the shortest-lived group. At day 0, immediately after eclosion, all groups started with similar levels of triacylglycerols, glucose, glycogen, and trehalose (Figure 4 C-F). Levels of TAGs, glucose and glycogen were significantly depleted in all groups after 4 days of starvation (Figure 4 C-E). Trehalose levels were slightly decreased in females housed in high temperature, long photoperiod condition but did not change in females housed in short photoperiod conditions at either temperature. This suggests that the starvation resistance observed in females housed in low temperature, short day conditions cannot be accounted for by changes in the utilization of TAGs, glucose, glycogen, or trehalose.

## DISCUSSION

We investigated maternal phenotypes associated with DP production in *Ae. albopictus* to identify those associated with temperature versus photoperiod. In general, we observed feeding and reproductive phenotypes that were influenced by temperature but were not predictive of DP egg status. While temperature influenced combined meal size in a two-choice assay between sugar meals with and without protein, neither blood nor sugar meal sizes were influenced by temperature or photoperiod in a no-choice context. Therefore, we did not find changes in maternal nutrient intake that predict DP egg status. We also report that temperature impacts egg retention and oocyte heterogeneity before blood feeding; however, these differences were not associated with DP egg status as females housed in low temperature, long photoperiod conditions also showed these changes even though they produce non-DP eggs. This suggests that differences in egg development pathways such as insulin signaling, lipid processing, and/or signaling by Juvenile Hormone or ecdysteroid (15,43–45) impact DP status by influencing features that are not detectable in overall maternal food intake or gross oocyte morphology.

Surprisingly, we found that housing adult females in short photoperiod, low temperature conditions were sufficient to increase starvation resistance. This may serve as a predictor of DP egg status in females before stress resistant eggs are produced. These findings also suggest that DP egg-producing females undergo metabolic changes before blood is consumed and egg production begins. Photoperiod has been found to have inconsistent impacts on adult body size. Costanzo and colleagues (46) found that *Ae. albopictus* females reared under short photoperiod conditions throughout larval development have larger body sizes, while Yee and colleagues (47) found that short photoperiod rearing decreased body size. We note that DP egg status was not evaluated in these studies and our protocol was designed to test changes after both larval feeding and female development were complete. Neither Ohtsu and colleagues (48) nor Yee and colleagues (46) observed any effect of photoperiod on lifespan when animals were provided *ad libitum* access to sugar. Similar to our findings (Figure 2G), Ohtsu and colleagues (48) report high levels of egg retention under short photoperiod conditions (12L:12D), although they found no evidence of tradeoffs between diapause incidence and reproduction or longevity. Our data suggests that metabolic changes and starvation resistance can be conferred by environmental exposure after larval development, suggesting metabolic plasticity in *Ae. albopictus* adult females.

One remaining question is the basis of the starvation resistance observed in our study. We observed that females housed in all three conditions started with similar levels of TAGs, glucose and glycogen, immediately after eclosion and that all of these nutrient stores were significantly depleted after 4 days of starvation. This means that the increased starvation resistance of females housed under low temperature, short photoperiod conditions cannot be accounted for by slow utilization or increased initial reserves of these nutrients. Prediapause *Drosophila* accumulate triacylglycerols, but not glycogen, and in many overwintering insects, triacylglycerols are the major fuel for basal metabolism (49). However these lipids are not always efficient since they become poor substrates for lipases upon solidification and the composition of saturated and unsaturated triacylglycerols differs with DP status in *Drosophila* (50,51). Females experiencing short photoperiods may undergo metabolic changes that alter the composition or ratios of lipid species that are not reflected in overall levels of TAG storage and could be studied using additional lipidomic and metabolomic analysis. Indeed, previous studies have identified differentially abundant lipids in DP eggs and have identified DP-specific metabolites that are similar to steroid derivatives (30), although it is not yet clear whether the similar maternal metabolic changes underlie the starvation resistance seen in our study.

This work identifies maternal phenotypes associated with temperature versus photoperiod. While many feeding and reproductive outputs were influenced by temperature in a way that did not predict DP egg status, unexpectedly, experiencing short photoperiod increases starvation resistance in *Ae. albopictus* females. Further exploration of the pathways that influence DP egg production and maternal metabolism may enable new targets to prevent *Ae. albopictus* from overwinter survival and provide new tools to limit range expansion of this invasive species.

## Acknowledgements

We thank the members of the Duvall lab and Dr. Oliver Hobert for comments on the manuscript, Jared Gatto and Dr. Mimi Shirasu-Hiza for technical assistance with TAG, glucose, glycogen, and trehalose assays.

## Author Contributions

IHL and LBD conceptualized the study. IHL, ASL, HVK and LD performed experiments and collected and analyzed data. LD and ASL designed and built the custom locomotor activity monitoring system. IHL and LBD wrote the manuscript with contributions from ASL, HVK, and LD. LBD provided guidance and supervision.

## METHODS

### Mosquito rearing and maintenance

*Aedes albopictus* (Foshan strain) were common-garden reared in environmental rooms (Harris Environmental) and/or incubators (Percival Scientific) at 80–90% relative humidity, 26°C, with a photoperiod of 16L:8D. Eggs were hatched in 500 mL of hatch broth (one crushed TetraMin fish food tablet in 1 L deionized, deoxygenated water). Larvae were fed crushed TetraMin fish food tablets. At the pupal stage, animals were split evenly to three environmental conditions and were housed in these conditions as adults (Figure 1A).

For assays involving unmated females, animals were separated by sex as pupae to ensure the unmated status of the females. For assays involving mated females, males and females were allowed to cohabit as adults for at least 4 days to ensure that females were mated. Adult mosquitoes were housed in BugDorm mesh cages with access to 10% sucrose and adult females were blood fed on defibrinated sheep blood (Hardy Diagnostics DSB100) using an artificial membrane feeder (Hemotek Ltd. SP6W1-3) for heated blood delivery (37°C). Whatman filter papers (Cytiva09805-F) lined in 8-oz plastic cups (VWR8900-664) half filled with DI water were placed in cages as an oviposition substrate.

### Female sexual maturity assay

Males and females were allowed to cohabit as adults for 4 days post-eclosion to ensure that females were mated. Blood feeding was carried out as described above and females were scored as “fed” based on visual inspection of blood in the midgut. Insemination status was determined by spermathecal dissection and scoring for spermathecal filling as described in (21).

### Hatching assay

We used a hatching assay adapted from Trpis, 1970 (17). Briefly, eggs were transferred to a 6-well mesh plate (CellTreat # 230605 PC 8.0µm insert). 2 mL of 1X PBS and 2 mL of 100% Clorox Germicidal Bleach (8.25% sodium hypochlorite—yields 7.85% available chlorine) were added to each experimental well using a disposable pipette to clear the cuticle to check for embryonation. The lids were replaced and covered with foil and rotated for 20 minutes in the dark. Eggs were then washed 4 times in 2 mL of 4°C 1X PBS for 10 minutes. Total number of eggs hatched was counted and embryonation status of the remaining unhatched eggs was determined by the presence or absence of eyes on either side of the egg. Hatching rates were determined by counting [(the total number of eggs hatched) / (total number of eggs hatched + embryonated unhatched eggs)] x 100.

### Oviposition assay

Blood-fed females were individually housed in 8-oz plastic cups (VWR8900-664) lined with Whatman filter paper and ∼25 mL of DI water for oviposition substrate. DI water was refilled as needed. Laid eggs were counted daily by eye for 14 days. Then, females were cold anesthetized at 4°C and their ovaries were dissected to count the number eggs retained using a Motic binocular microscope.

### Egg size measurement

Laid eggs were dried on flat Whatman filter paper for 2 - 3 days and imaged directly using a Motic binocular scope. Images of eggs were taken at 4X magnification. Egg width and length measurements were taken along the longest perpendicular axes of eggs and converted to μm using Motic Images Plus 3.0 software.

### Two-choice assay

Females were provided access to 10% sucrose wick on the first day after eclosion and then starved for 3 days with access to only DI water. The assay was set up in an 8-oz plastic cup (VWR8900- 664) covered with mesh, with two 50 mLFalcon tube caps containing 7 mL each of either 10% sucrose or 10% bovine serum albumin (BSA) in 10% sucrose solution. Each meal was labeled with either 1% rhodamine B (absorption/emission: 540 nm/575 nm) (Sigma-Aldrich R6626) or 0.002% fluorescein (487 nm/528 nm) (Sigma-Aldrich F6377). 8–10 animals were put into each plastic cup and placed into the corresponding environmental conditions (Figure 1A) for 24 hours. After 24 hours, females were cold-anesthetized, individually collected in sterile 1.5 mL microcentrifuge tubes, and stored at -20°C until ready to measure. Fluorescence measurements were taken on a Biotek Synergy Plate Reader on a 96-well glass bottom plate (ThermoScientific 152037). The samples were each homogenized in 100 μL of 1X PBS and centrifuged to isolate supernatant. 50 μL of the supernatant of each sample was transferred to a 96-well plate. Samples were run alongside a standard of 2-day starved females homogenized in 1X PBS with known concentrations of fluorescein or rhodamine B.

Analysis of the fluorescence was performed on Microsoft Excel using known standards. Preference index was calculated as (%BSA+Sucrose meal - % Sucrose meal).

### No choice sugar feeding assay

Adult females were starved for 24-48 hours prior to blood feeding and provided access only to DI water. The assay was set up in an 8-oz plastic cup (VWR8900-664) covered with mesh, with a 50- mL Falcon tube cap containing 7 mL each of 10% sucrose with 0.002% fluorescein. 8–10 animals were put into each plastic cup and placed into the corresponding environmental conditions and allowed to feed *ab libitum* for 2 hours. After 2 hours, females were collected and meal size was quantified using fluorescence readings taken on a BioTek Synergy Plate reader as described in the “Two-choice assay” section.

### No choice blood feeding assay

Pre-blood fed females were starved 48 hours prior to blood feeding and provided access only to DI water, cold anesthetized and weighed in groups of 3. Once females awoke from anesthetization, same females were blood fed to repletion using methods described in “mosquito rearing and maintenance” and then re-weighed individually. Blood consumption was calculated as (weight of post-blood fed female - weight of pre-blood fed female).

### Ovarian development scoring

Ovaries were dissected before blood feeding and 1, 3, 7, and 14 days after a blood meal. Females were cold anesthetized on ice and fixed in 4% PFA (ThermoScientific 28906) for 2 hours. Using a Motic binocular stereo microscope, the head, wings, and legs were removed with dissection forceps. A droplet of DI water was placed on the dissection arena to prevent desiccation of the ovaries. The ovaries were then dissected by pulling down from the last segment of the abdomen into a new droplet of water. Pairs of ovaries were mounted in Vectashield (Vector Laboratories H-1000-10) and imaged on a Nikon Ti2E microscope. Ovarian developmental stage was determined by measuring the proportion of the ovariole occupied by the nurse cells using Nikon NIS-elements software.

### Starvation resistance

Females were collected immediately after eclosion and provided *ad libitum* access to DI water wicks to prevent desiccation. The starvation resistance assay was carried out under the same housing conditions shown in Figure 1A. Every 24 hours, the total number of dead animals were counted until no animals remained.

### Locomotor activity assay

Three identical “MozzieMonitor” locomotor activity assays were constructed and placed under three environmental conditions (Fig 1A) to record daily activity level. The design of the assay is similar to Flybox, which uses an IR camera platform to record insect locomotion (22). Briefly, monitors consisted of RaspberryPi Camera Module 3 NoIR Wide cameras mounted on top of the acrylic frames. The Pi cameras were controlled by a Python (version 3.9.2) script installed on Raspberry Pi 4 B 4 GB computers to take an image every minute. Sets of 850nm IR LED strips (Waveform Lighting 7041.850.5M) were installed on the bottom of the frames to provide constant IR light for videography. Two cooling fans (Newegg 9SIB7W5HBZ4935) were installed underneath each set of IR LED strips to dissipate heat.

To measure locomotor activity, mosquitoes were housed individually in the wells of 6-well tissue culture plates (CellTreat 229106). Each well was supplied with 3 mL of 0.5% agarose gel (IBI Scientific IB70050) to provide humidity and water source. The 6-well plates were inverted in the MozzieMonitor to avoid water condensation. A piece of Kimwipe (Sigma-Aldrich Z188956) was inserted between the cover and the base of each 6-well plate to provide a rough and absorbent surface for mosquitoes to walk on. Hot glue was used to tightly seal the 6-well plates.

Animals eclosed within 24 hours were loaded into 6-well plates at ZT8. Images of mosquitoes in 6-well plates were acquired starting from ZT16 of the same ZT day for 6 consecutive days. At the end of each experiment, images taken every minute were compiled as a 30 frames-per-second video using custom Matlab code (Version 2020b). The videos were analyzed with EthoVision (Version 17) where individual mosquitoes were tracked using dynamic background subtraction. Distances traveled between frames were extracted from EthoVision output files using a custom Python (version 3.10.12) code. All locomotor activity assays were performed 3 independent times with at least 12 animals for each condition. Animals that survived past ZT24 of the fifth recording day were included in the calculation of total activity from days 1 – 3 to avoid scoring activity of animals near death. Vector files used for laser-cutting, schematics of the MozzieMonitor components, and all codes used for the MozzieMonitor assay are deposited at [https://github.com/DongLinhan/MozzieMonitor].

### Triacylglyceride measurement

Samples were prepared from adult mosquitoes collected at 0 and 4 days post eclosion (dpe) from 3 environmental conditions (Fig 1A). As adults, mosquitoes were provided access to DI water only. Thin-layer chromatography (TLC) and image quantification was performed using published methods (23). Briefly, each biological replicate was made up of 5 age-matched mosquitoes that were placed in a 1.5 mL microcentrifuge tube and weighed. Samples were stored at -80°C until extraction. Samples were solubilized by homogenizing each sample in 250 µL of a 2:1 chloroform:ethanol mixture and centrifuged for 13 minutes at 12,000 rpm at 4°C. Supernatant was diluted four fold in the 2:1 chloroform:ethanol mixture and 10 µL of this diluted sample spotted on a silica gel, 5 μL at a time, with brief air-drying in between. (Sigma Aldrich Silica gel on TLC- PET foils 99577-25EA). Each plate was loaded with two age-matched conditions, with 4 biological replicates per condition. A reference standard of 100 sugar-fed female mosquitoes 3-5 dpe diluted 1:8 was included on each plate and run in triplicate. The TLC plates were developed with an eluent composed of 70 mL n-hexane, 30 mL diethyl ether, and 1 mL acetic acid. Plates were washed with 1 M NaCl for 10 seconds and then stained for 5 hours using 0.2% amido black (Sigma Product N- 3393) in 1 M NaCl. Amido black was then washed 2x with 1 M NaCl and 1x with DI water before being allowed to dry overnight. Plates were imaged on the colorimetric setting of a Bio-Rad GelDoc Go Imaging System. Images were quantified using FIJI, with the background of the plate subtracted from each sample and divided by the mean intensity of the reference standard. All relative TAG measurements were divided by the mass of the sample measured prior to lipid extraction.

### Carbohydrate analysis

Samples were prepared from adult mosquitoes collected at 0 and 4 days post eclosion with *ad libitum* access to water only. Carbohydrate quantification was performed using methods adapted from published protocols (23). Briefly, groups of 5 age-matched, *ad libitum* water fed female mosquitoes from each condition were measured and stored at -80°C until extraction. Samples were homogenized for 30 seconds in 200 µL of 1x PBS. Samples were then centrifuged at 14000 rpm at 4°C for 3 minutes. Supernatant was removed and heated at 70°C for 10 minutes before being centrifuged again at 14000 rpm at 4°C for 3 minutes. Supernatant was then removed and diluted 1:2 in 1x PBS. Samples were loaded onto clear, 96-well plates (Non-Treated Multiple Well Plates, Cell Treat 229597). Glucose was detected using the Glucose (HK) Assay kit (Sigma GAHK20) after incubation for 1 hour at 37°C. Glycogen was measured with the same kit using amyloglucosidase (Amyloglucosidase from *Aspergillus niger,* A1602 Sigma) digestion for 1 hour at 37°C. To measure trehalose, samples were prepared as above, but were diluted 1:2 in trehalase buffer (5 mM Tris, 137 mM NaCl, and 2.7 mM KCl, pH corrected to 6.6). Samples were then diluted 1:2 once more in trehalase digestion (Trehalase from porcine kidney, Sigma T8778), or trehalose buffer for control, before an 18-hour incubation at 37°C of both samples and standards. After the 18-hour incubation, samples were loaded into the 96-well plate and incubated with glucose reagent 37°C for 1 hour. Samples for glucose, glycogen and trehalose readings were run alongside standard curves with a 1:2 serial dilution of known concentrations of glucose (included in Glucose (HK) Assay kit), glycogen (Sigma-Aldrich G8876), and trehalose (D-(+)-Trehalose dihydrate from Saccharomyces cerevisiae, >99%, Sigma-Aldrich T9531). All reactions were stopped after the 1-hour 37°C incubation by adding 6 M sulfuric acid. A BioTek Synergy Neo2 Hybrid Multimode Reader was used to measure absorbance values at 540 nm. Glycogen and trehalose were measured on plates alongside glucose, and glucose values were subtracted from glycogen and trehalose absorbance readings. Glucose, glycogen, and trehalose measurements of samples were determined from known concentrations from the standard curves and then divided by the mass of each sample collected prior to the assay. The same samples were used across all three carbohydrate analyses.

### Quantification and Statistical Analysis

Statistical analysis was performed in Microsoft Excel (version 16.81) and GraphPad Prism (version 9.5.1). Data are shown as median with range or mean with SEM as indicated in the figure legends. One-way ANOVA with Tukey’s multiple comparison was used to compare more than 2 groups of normally distributed datasets, Kruskal-Wallis test with Dunn’s multiple comparisons test was used to compare more than 2 groups for non-normally distributed datasets, and Mann-Whitney tests were used for pairwise comparisons of non-normally distributed groups. Details of statistical methods are reported in the figure legends.

### Lead contact

Further information and requests for resources and reagents should be directed to and will be fulfilled by the lead contact, Laura B. Duvall

### Data and code availability

All data reported in this paper are available in Data File S1 and original code is publicly available via Github (https://github.com/DongLinhan/MozzieMonitor)

---

**Supplemental Figure 1.**
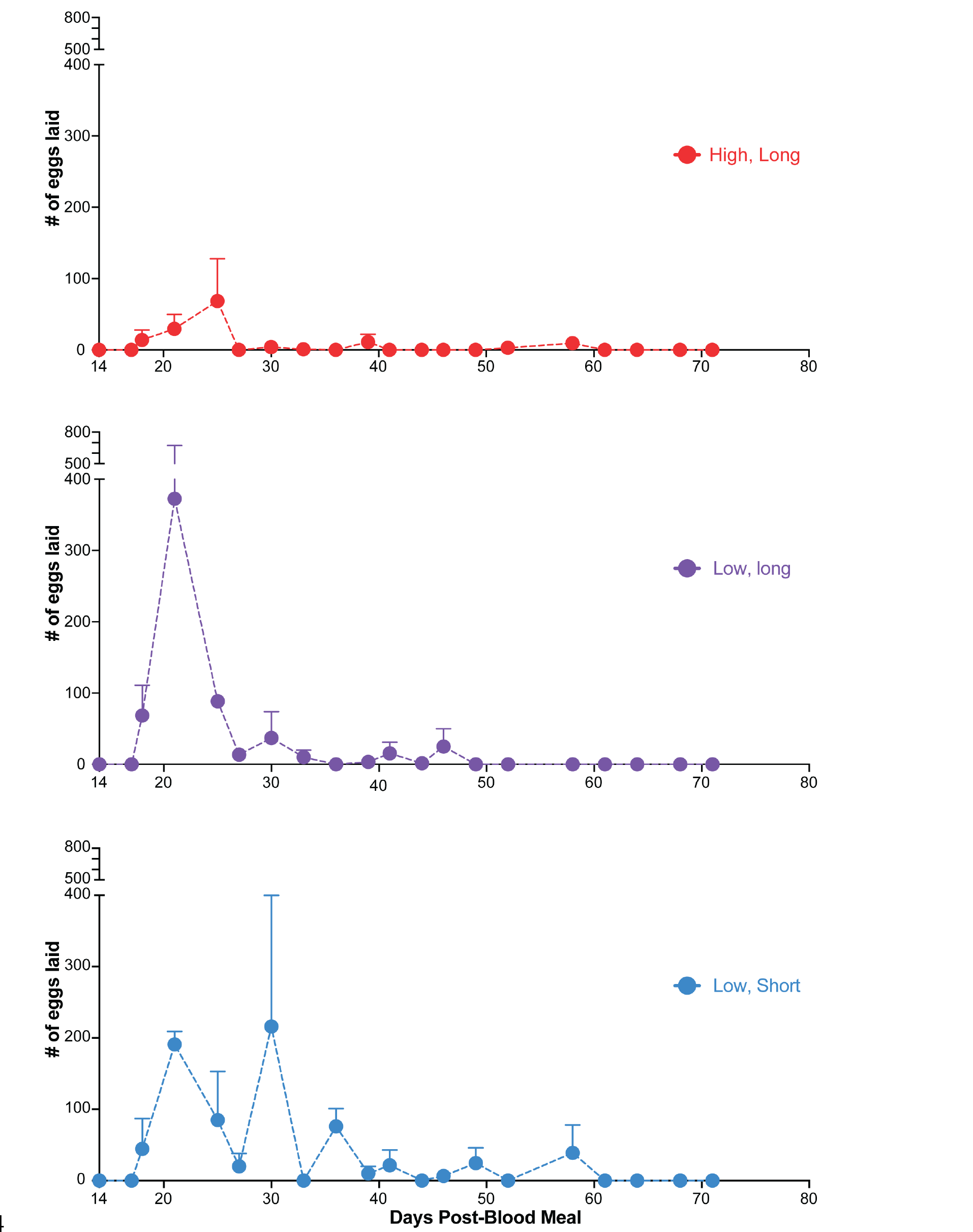
Females continue to lay retained eggs until death After a single blood meal, oviposition was allowed until death (data is shown as mean with SEM; n = 2 replicates; >23 females per replicate).

**Supplemental Table 1.** Female sexual maturation at 3 days post eclosion Adult females were allowed to mate freely with under three rearing conditions. Female sexual maturation was measured by both the filled spermathecae and ability to blood feed (n=11-15 females).

| Environmental Conditions | Spermathecae Filled | Blood-Fed |
| --- | --- | --- |
| High, Long | 100% | 87% |
| Low, Long | 91% | 100% |
| Low, Cool | 92% | 75% |

## Notes

https://github.com/DongLinhan/MozzieMonitor

### Competing Interest Statement

The authors have declared no competing interest.

